# A proteolytic AAA+ machine poised to unfold a protein substrate

**DOI:** 10.1101/2023.12.14.571662

**Authors:** Alireza Ghanbarpour, Robert T. Sauer, Joseph H. Davis

## Abstract

AAA+ proteolytic machines unfold proteins prior to degradation. Cryo-EM of a ClpXP-substrate complex reveals a postulated but heretofore unseen intermediate in substrate unfolding/degradation. The natively folded substrate is drawn tightly against the ClpX channel by interactions between axial pore loops and the substrate degron tail, and by contacts with the native substrate that are, in part, enabled by movement of one ClpX subunit out of the typically observed hexameric spiral.

## MAIN TEXT

From bacteria to mammals, ATP-fueled AAA+ proteases degrade regulatory, unneeded, or damaged intracellular proteins (Sauer and Baker 2011). For target proteins with stable three-dimensional structures, ClpXP and other AAA+ proteases harness the energy of ATP hydrolysis to unfold this structure before translocating the denatured polypeptide through a narrow axial channel and into a self-compartmentalized peptidase chamber for degradation. Within the ClpXP complex, the ClpX unfoldase mediates substrate specificity by recognizing short ‘*degron*’ peptides within target substrates. Unfolding is thought to occur when ClpX translocates a disordered substrate segment, containing a recognition degron, until the attached native structure is pulled against the axial channel. Subsequent power strokes then attempt to further translocate the substrate polypeptide through the channel, resulting in repeated application of a force that increases the cumulative probability of unfolding and eventually succeeds in denaturing the substrate to allow sequence-independent translocation of the denatured polypeptide into the ClpP peptidase chamber for proteolysis (**Fig. 1a**). This model accounts for many studies (Sauer, Fei *et al*. 2022). However, visualization of a native substrate being pulled against the axial channel of any AAA+ protease has not, until now, been achieved.

**Figure 1:**
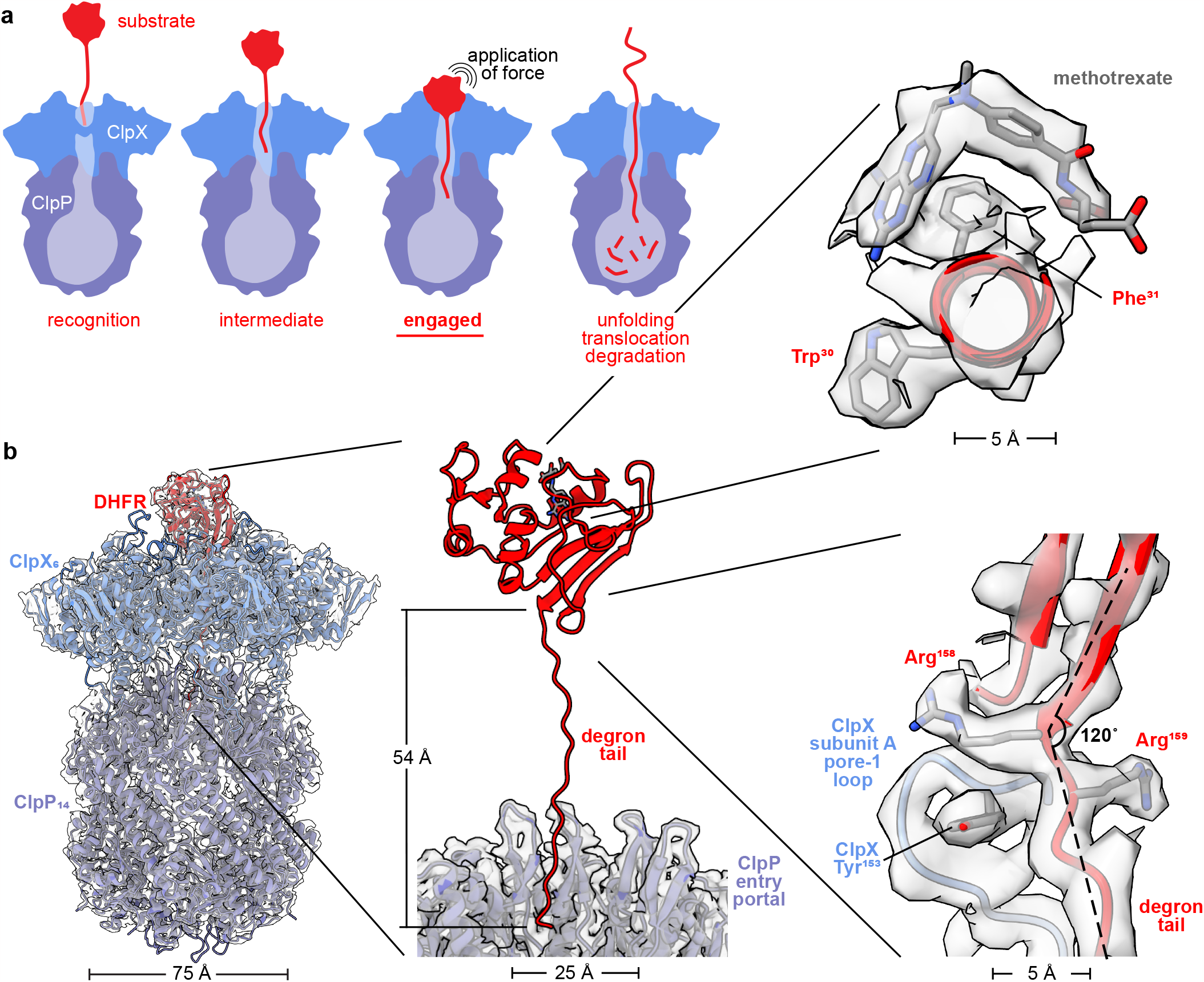
ClpXP bound to degron-tagged DHFR. **(a)** Cartoon of steps in ClpXP degradation of a protein substrate. **(b)** Overlay of cryo-EM map and model of ClpXP bound to the degron-tagged DHFR substrate (left). Density for the degron tail could be modeled extending from the folded domain of DHFR to the ClpP entry portal (center). The insets (right) display density for methotrexate and the DHFR Trp^30^ and Phe^31^ side chains (top), and the last two residues of native DHFR, Arg^158^, and Arg^159^, contacted by the pore-1 loop of ClpX subunit A, which includes Tyr^153^.

To visualize this postulated ‘*engaged*’ complex, we prepared cryo-EM grids using *Escherichia coli* ClpX^ΔN^, a variant lacking the N-terminal domain, *E. coli* ClpP, and a protein substrate consisting of *E. coli* dihydrofolate reductase (DHFR) with a C-terminal tail containing a ssrA degron (see **Methods**). Along with ATP, the reaction mix included methotrexate (MTX), a small-molecule that binds to and stabilizes DHFR (Ainavarapu, Li *et al*. 2005), and prevents its degradation by ClpXP (Lee, Schwartz *et al*. 2001). After grid screening, data collection, and image processing (see **Methods; Figs. S1-S4**), we determined a structure at ~2.8 Å resolution (**Table 1; Fig. 1b**), which showed natively folded DHFR in close contact with the ClpX ring and 19 residues of the DHFR degron tail in a β-ribbon conformation that extended ~50 Å through the axial channel of ClpX and into the entry portal of ClpP.

**Table 1:**
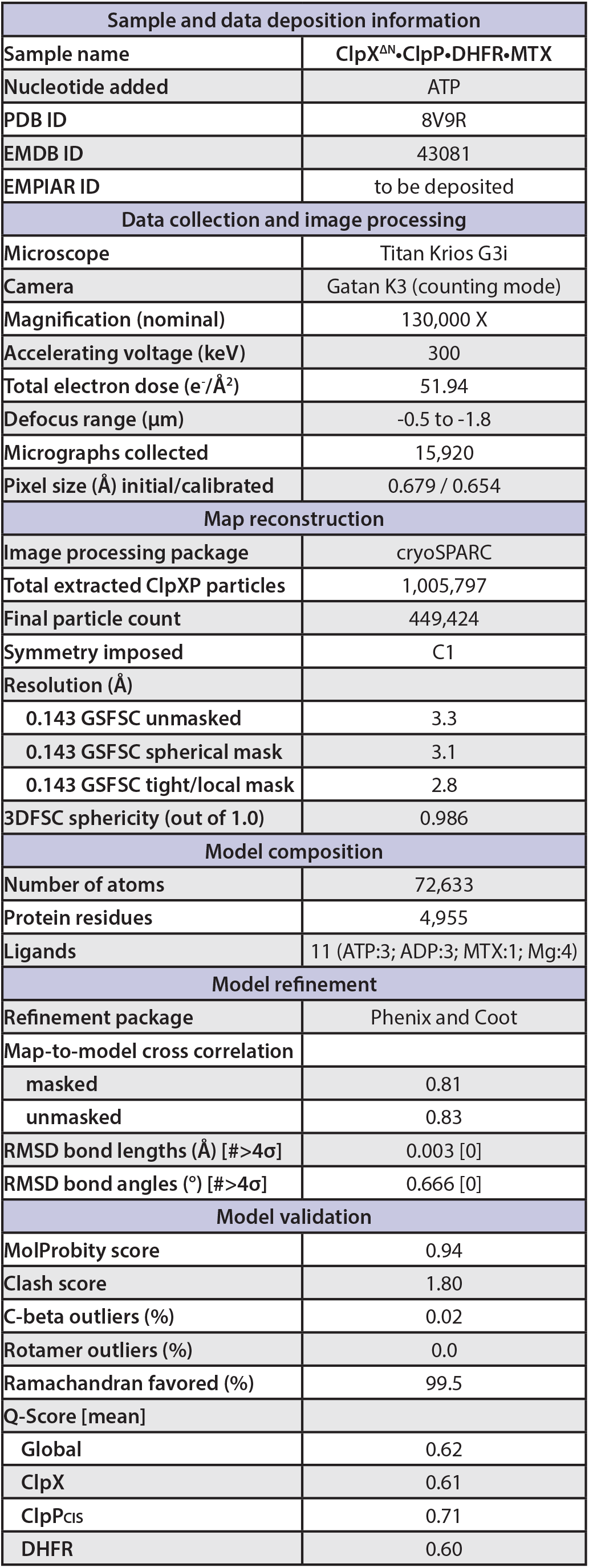
Cryo-EM data collection, processing, model building, and validation statistics.

Density for DHFR and MTX was well resolved in the map (**Fig. 1b**), and the refined model aligned to a DHFR•MTX crystal structure (Sawaya and Kraut 1997) with a Cα RMSD of 1.25 Å. Thus, ClpX binding *per se* does not induce partial denaturation of DHFR. Indeed, ClpX supports ClpP degradation of a very large number of structurally diverse proteins (Roche and Sauer 2001, Flynn, Neher *et al*. 2003, Neher, Villen *et al*. 2006), making it unlikely that an ability to selectively bind partially denatured conformations of any particular protein would have evolved.

As newly highlighted in our structure, packing between the native DHFR domain and ClpX determines the angle between the degron tail and terminal DHFR structural element (**Figs. 1b, 2a**). The C-terminal β-strand of native DHFR, which is embedded in a β-sheet, connected to the degron tail at an angle of ~120° and without slack (**Fig. 1b**). Arg^158^, the penultimate residue of untagged DHFR, had the same conformation as in an MTX-bound crystal structure (Sawaya and Kraut 1997), whereas the side chain of Arg^159^, the last native residue, assumed a new conformation that allowed the top axial pore-1 loop of ClpX to contact the extreme C-terminus of the native DHFR domain (**Fig. 1b**). We posit that the absence of slack would ensure that force from a power stroke is directly applied to the native substrate. We expect that the observed contact angle between substrate and unfoldase also likely influences unfolding efficiency, as some angles would optimize peeling of secondary-structure elements, whereas others would require shearing of multiple hydrogen bonds (Rohs, Etchebest *et al*. 1999).

Grip between the axial-channel loops of ClpX and a degron tail also affects unfolding efficiency, with larger non-polar side chains in the tail providing superior grip and faster unfolding (Sauer and Baker 2011, Bell, Baker *et al*. 2019, Sauer, Fei *et al*. 2022). In our structure, five pore-1 loops (subunits A,B,C,D,E) and four pore-2 loops (subunits A,B,C,D) of ClpX_6_ contacted 10 DHFR-degron residues in the channel (**Fig. 2a**), with many interactions involving large side chains of the degron tail. The pore-1 loop of subunit F was disengaged from the degron tail but close to native DHFR. Hundreds of unsuccessful ClpX power strokes can be needed to unfold and degrade a stable substrate (Kenniston, Baker *et al*. 2003, Cordova, Olivares *et al*. 2014). In this regard, the extensive contacts between ClpX and the degron tail are likely to minimize substrate dissociation after an unsuccessful power stroke, thereby increasing the cumulative probability of unfolding for MTX-free DHFR. Our structure also resolved a peripheral collar of contacts between the six RKH loops of ClpX_6_ and native DHFR (**Fig. 2b**), which would also minimize substrate dissociation during denaturation attempts. Moreover, we expect that the marked flexibility of individual RKH loops observed in this and previous structures (**Fig. S5**) allows ClpX to accommodate the highly divergent structures of the very large number of substrates it degrades. We compared Cα positions of Arg^228^ in the RKH loops of our DHFR complex with the corresponding positions in an ssrA-degron complex (Fei, Bell *et al*. 2020) (pdb 6WRF) and found movements ranging from 12 to 30 Å (**Fig. S5)**.

**Figure 2:**
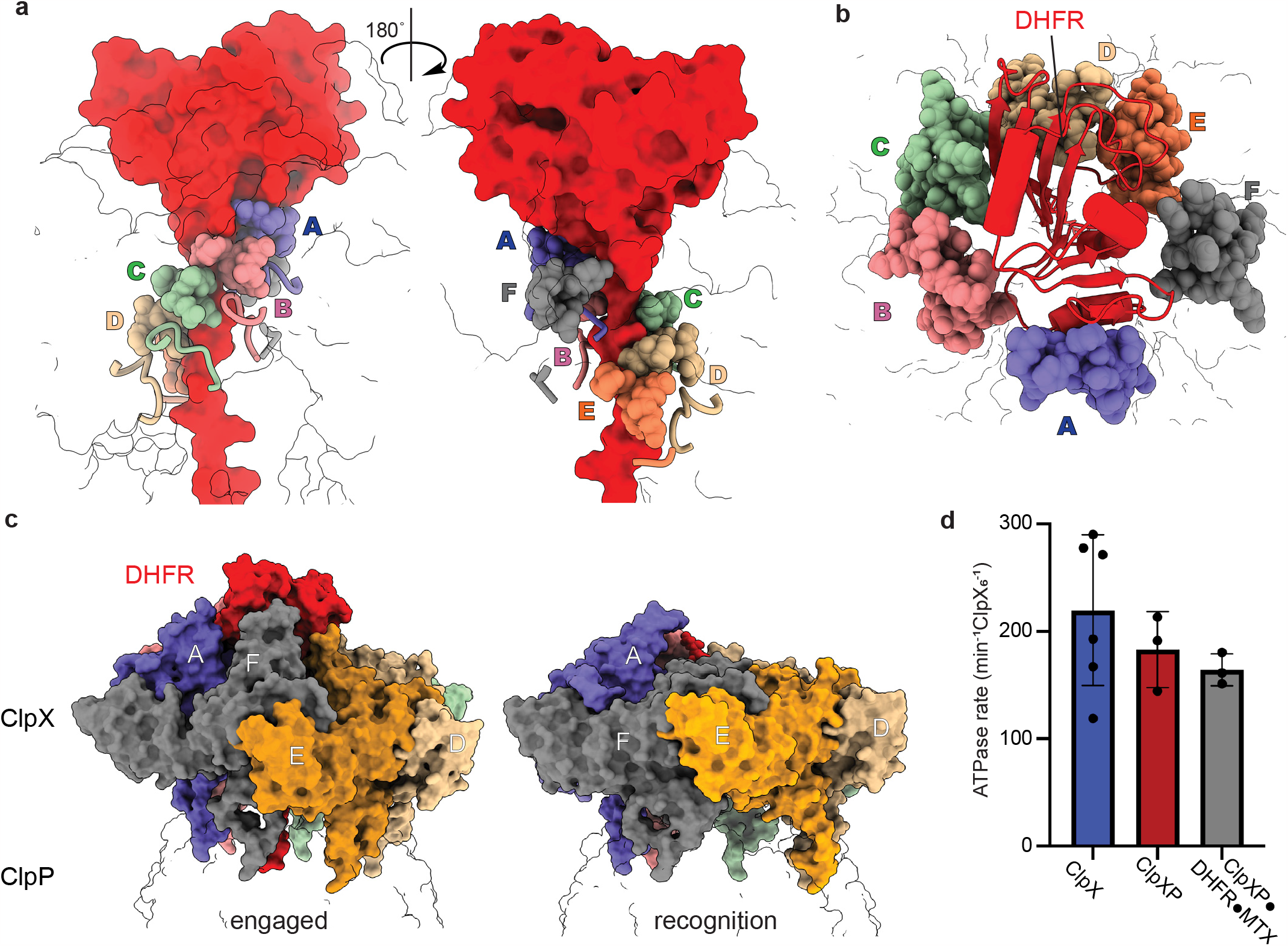
ClpX•DHFR interactions and the consequences of binding. **(a)** Front and back views of contacts between pore-1 loops of ClpX (residues 150-155; sphere representation), pore-2 loops of ClpX (residues 198-205; cartoon representation), and DHFR and its degron tail (red surface representation). The six ClpX subunits are annotated A-F and colored purple, salmon, green, wheat, orange, and grey, respectively. The ClpX subunit F pore-1 loop is disengaged from the degron tail but adjacent to the folded domain of DHFR. **(b)** Top view showing packing of the ClpX RKH loops (residues 220-238; sphere representation colored by subunit as above) around native DHFR (red cartoon representation). **(c)** Side views of ClpX complexes with DHFR (left) and an ssrA degron (right; pdb 6WRF) illustrating major upward movement of subunit F. **(d)** Bars show mean rates of ATP hydrolysis for ClpX, ClpXP, or ClpXP in the presence of degron tagged DHFR•MTX ± 1 SD, with symbols denoting replicate assays.

In most ClpX hexamers, the axial pore-1 loops form a spiral with the A loop at or near the top and the F loop near the bottom (Fei, Bell *et al*. 2020, Fei, Bell *et al*. 2020, Ripstein, Vahidi *et al*. 2020, Ghanbarpour, Cohen *et al*. 2023, Ghanbarpour, Fei *et al*. 2023). By contrast, subunit F moved ‘up’ in our DHFR-bound structure **(Fig. 2c)**, resulting in its pore-1 loop being ~20 Å higher in the channel than the pore-1 loop of subunit E, disengaged from the DHFR degron tail, and at roughly the same level as the pore-1 loop of subunit B (**Fig. 2a**). Only one other ClpX_6_ structure, which was engaged with the SspB adaptor and a GFP substrate (Ghanbarpour, Fei *et al*. 2023), has a similar ‘up’ conformation of subunit F. These engaged structures may be specialized for substrate denaturation. In both the DHFR- and SspB/GFP-bound structures, the ClpP-binding IGF loop of subunit E also hopped into a cleft in the ClpP_7_ ring that is unoccupied in subunit-F ‘down’ structures (**Fig. S6**).

DHFR-bound ClpX had ATP bound in subunits A/B/C and ADP in subunits D/E/F (**Table S1; Fig. S7**), whereas other ClpX_6_ structures (Fei, Bell *et al*. 2020, Fei, Bell *et al*. 2020, Ripstein, Vahidi *et al*. 2020, Ghanbarpour, Cohen *et al*. 2023, Ghanbarpour, Fei *et al*. 2023) contain four or five ATP or ATPγS molecules and one or two ADPs. We considered the possibility that DHFR•MTX binding might trap ClpXP in an ATPase inactive conformation by keeping subunit F in an ‘up’ conformation, but the complex hydrolyzed ATP at a rate only slightly slower than ClpX or ClpXP alone (**Fig. 2d**). We also used cryoDRGN (Zhong, Bepler *et al*. 2021, Kinman, Powell *et al*. 2023, Sun, Kinman *et al*. 2023) to test for minor populations of ClpX or DHFR structures in the complex, specifically looking for structures in which subunit F was ‘down’ or the IGF-loop of subunit E ‘hopped’, as such conformers were expected had ClpX successfully unfolded DHFR and begun translocating the denatured polypeptide. No such state was observed and, instead, we observed DHFR moving subtly or becoming unresolved above the pore, possibly as a result of ATP-hydrolysis-dependent power strokes within ClpX (**Movie S1**). Combined with our ATPase assay, this dynamic view of the structure suggests that ClpX readily hydrolyzes ATP as it attempts to denature DHFR•MTX, even when this outcome is highly improbable.

## MATERIALS AND METHODS

### Protein purification

*E. coli* ClpP-His_6_ and a C-terminally His_6_-tagged ClpX^ΔN^ variant consisting of three ClpX^ΔN^ subunits connected by genetically encoded peptide linkers were expressed separately in *E. coli* and purified by Ni^++^-NTA, ion-exchange, and gel-filtration chromatography (Martin, Baker *et al*. 2005). The *E. coli* DHFR gene was fused to the GSHLGLIEVEKPLYCVEPFVGETAHFEIELSEPDVHGQWKLTSH_6_ peptide tail at its C-terminus and cloned into a pETDuet expression vector. This tagged DHFR protein was expressed in BL21 (DE3) cells at 18 °C overnight and purified using Ni^++^-NTA affinity chromatography by loading and washing in buffer N1 [20 mM HEPES pH 7.8, 400 mM NaCl, 100 mM KCl, 10% glycerol, 1 mM DTT, 15 mM imidazole] and eluting in buffer N1 supplemented with imidazole to a concentration of 250 mM. The eluate was concentrated and then subjected to gel-filtration chromatography in buffer GF [20 mM HEPES pH 7.5, 300 mM KCl, 10% glycerol, and 1 mM TCEP]. A synthetic degron-tag peptide consisting of maleimide-GSGSWSHPQFEK<u>AANDENYALAA</u> (21st Century Biochemicals, Inc.), where the underlined sequence is the ssrA tag, was crosslinked to the tagged DHFR protein by reacting ~20 equivalents of the peptide with one equivalent of tagged DHFR for 2 h at room temperature in buffer CL [100 mM HEPES, pH 7.5, 500 mM NaCl, 10% glycerol, 1 mM TCEP] that had been degassed using argon. Unreacted peptide was quenched using 20 mM DTT, removed using a PD-10 desalting column, and the crosslinked DHFR-C15-ssrA protein was concentrated and flash-frozen in liquid nitrogen for storage. We used this branched-tail substrate hoping to visualize how ClpX accommodates multiple polypeptides in its axial channel, but the branch was not well ordered and could not be modeled in the structure.

### Cryo-EM single particle analysis

The ClpX^ΔN^ variant (5.7 μM pseudohexamer) and ClpP (1.5 μM tetradecamer) were incubated with ATP (5 mM) and DHFR-C15-ssrA (20 μM) in buffer R1 [20 mM HEPES pH 7.5, 100 mM KCl, and 25 mM MgCl_2_] at room temperature for 5 min. Prior to vitrification, 2.5 μL sample aliquots were placed on 200-mesh Quantifoil 2/1 copper grids, which had been glow-discharged at –15 mA for 60 s using an easiGlow discharger (Pelco), and samples were blotted using a FEI Vitrobot Mk IV with a 0 blot force at 6 °C and 95% relative humidity for 4 s.

Movies were collected with EPU (Thermo Fisher Scientific) using aberration-free image shift (AFIS) and hole-clustering methods on a Titan Krios G3i (Thermo Fisher Scientific) with an acceleration voltage of 300 keV and magnification of 130,000 X and detected in super-resolution mode on a K3 detector (Gatan) with a pixel size of 0.679 Å (binned by 2). Movies were collected as 40 frames with a total exposure per specimen of 51.94 e–/Å^2^ and a defocus range from –0.5 to –1.8 μm. Data processing was performed in cryoSPARC (v.3.3.1) (Punjani, Rubinstein *et al*. 2017) as depicted in **Figure S1**. Raw movies (15,290) were pre-processed using ‘patch motion correction’, and ‘patch CTF estimation’. Particles (~47,000) picked using the blob-picker tool from 1,000 random micrographs were extracted (box size 440 px, Fourier cropped to 256 px) and subjected to ‘2D classification’. A set of 4 well-resolved 2D classes composed of side and shoulder views were provided to the ‘template picker’ tool and applied to the full dataset. The particles and micrographs were subjected to ‘manually curate exposures’, filtering out low resolution and low particle count exposures. Particles were extracted (box size 440 px, Fourier cropped to 256 px) and after two rounds of 2D classification, the preliminary stack included 1,005,797 particles.

Multiclass ‘ab-initio’ reconstruction was performed using three classes. One class consisted of 629,143 ClpXP particles (group I) and another of 251,567 ClpXP particles (group II). Inspection of the remaining class via 2D classification revealed a mixture of free ClpP, a low-resolution ClpXP complex, and ‘junk’ particles likely corresponding to errantly picked particles, and these particles were not further considered. After separate homogeneous refinements of groups I and II, heterogeneous refinement was performed using six sub-classes of group I and four sub-classes of groups II. The resulting maps were visually inspected, and four classes from group I and three classes from group

II were selected for another round of homogeneous refinement (801,567 particles) that used an ab initio map from group I as an initial model. These aligned particles were then recentered on ClpX using ‘volume tools’ followed by homogeneous refinement with per-particle defocus estimation enabled and subjected to another round of heterogeneous refinement (four classes) that identified the 449,424 particles used for final reconstructions. The final map was obtained through homogeneous refinement, followed by local refinement employing a mask focused on ClpX and the cis ClpP ring. The final map was rescaled using a calibrated pixel size of 0.654 Å (Ghanbarpour, Cohen *et al*. 2023) in cryoSPARC and had a GSFSC of ~2.8 Å after FSC-mask auto-tightening. After centering particles on the ClpP equatorial ring and extracting the centered particles with a larger box size, we also visualized a second ClpX•DHFR complex bound to the second heptameric ring of ClpP_14_, albeit at lower resolution (**Fig. S8**).

Local resolution was estimated using MonoRes (Vilas, Gomez-Blanco *et al*. 2018) within cryoSPARC; angular FSCs were calculated using the 3DFSC server (Tan, Baldwin *et al*. 2017); and Q-scores were calculated using a ChimeraX Q-score plugin (Pintilie, Zhang *et al*. 2020). Model building was performed using a combination of ChimeraX-1.3 (Pettersen, Goddard *et al*. 2021), Coot-0.9.4 (Casanal, Lohkamp *et al*. 2020), and Phenix -1.14 (Liebschner, Afonine *et al*. 2019). The final map was sharpened using CryoSPARC with a b-factor of -50.

### ATPase assays

ATP hydrolysis was measured using a coupled enzymatic reaction (Norby 1988) in which NADH oxidation to NAD+ reduces absorbance at 340 nm (Δε = 6.22 mM^-1^ cm^-1^) using a SpectraMax M5 plate reader and a 384-well assay plate (Corning, 3575). A stock ATPase reaction mix (20X) contained 20 μL of a mixture of pyruvate kinase and lactic dehydrogenase from rabbit muscle (P0294, Sigma Aldrich), 10 μL of 200 mM NADH (CAS# 606688), 15 μL of 1 M phosphoenolpyruvate (Sigma Aldrich, 10108294001) in 25 mM HEPES-KOH (pH 7.6), and 25 μL of 200 mM ATP (pH 6.5).

For assays, ClpX_6_ (1 μM) with or without ClpP_14_ (3 μM) and DHFR-`GSYLAALAA` (Bell 2020) plus MTX (16 μM each) was present in 10 μL of buffer AB [25 mM HEPES pH 7.8, 100 mM KCl, 20 mM MgCl_2_, 10% glycerol]. After 5 min of incubation at 30 °C with 2 mM ATP, the ATPase assay was initiated by addition of an equal volume of 2X ATPase reaction mix in buffer AB. Final reaction concentrations were: 0.5 μM ClpX_6_, 1.5 μM ClpP_14_ (if present), 8 μM DHFR-`GSYLAALAA` with MTX (if present), ATPase reaction mix (1X) in a total reaction volume of 20 μL.

### CryoDRGN analysis

CryoDRGN was used to analyze the full set of 449,424 particles where the signal for the trans ring of ClpP had been subtracted in cryoSPARC by: 1) aligning particles on ClpP_14_ through a local refinement with a mask surrounding ClpP_14_; 2) subtracting signal of the trans ClpP_7_ ring; and 3) performing a final local refinement using the signal subtracted particle stack and a mask encompassing DHFR, ClpX_6_, and the ClpP_7_ cis ring. These particles were down-sampled to a box size of 254 px (1.13 Å/pixel) and used to train an eight-dimensional latent-variable model in cryoDRGN v2.3.0 using 1024x3 encoder and decoder architectures. The poses and ctf parameters for cryoDRGN training were supplied from the aforementioned local refinement. Following 20 epochs of training, 100 volumes were sampled from the k-means cluster centers of latent embeddings. After extensive visual inspection of these volumes using our atomic model as a reference, we observed neither the ‘down’ conformation in F subunit nor any instances of IGF ‘hopping’. However, the sampled volumes did reveal subtle movements of DHFR or instances where it was unresolved (**Supplemental Movie 1**).

## ACKNOWLEDGMENTS

We thank Ed Brignole, Laurel Kinman, and Barrett Powell for helpful discussion and feedback, and the MIT-IBM Satori team and the MIT SuperCloud Supercomputing Center for high performance computing resources and support. This work was supported by NIH grants R01-GM144542 and R35-GM141517, and NSF-CAREER grant 2046778. Samples were prepared at the Automated Cryogenic Electron Microscopy Facility in MIT.nano and screened on a Talos Arctica microscope, which was a gift from the Arnold and Mabel Beckman Foundation.

## CONFLICTS OF INTEREST

The authors declare no conflicts of interest.

## CONTRIBUTIONS

AG purified proteins, performed biochemical assays, prepared samples for EM imaging and collected data. AG and JHD processed EM data and performed reconstruction and refinement. AG and RTS built and refined the atomic models. All authors contributed to writing and editing the manuscript. JHD and RTS supervised the project.

## SUPPLEMENTARY TABLE AND FIGURES

**Table S1.** Nucleotide occupancy. Nucleotide occupancy of subunits in different ClpX cryo-EM structures (Fei, Bell *et al*. 2020, Fei, Bell *et al*. 2020, Ripstein, Vahidi *et al*. 2020, Ghanbarpour, Cohen *et al*. 2023, Ghanbarpour, Fei *et al*. 2023). Red designates subunits that appear catalytically active for ATP hydrolysis. Blue designates subunits that are catalytically inactive either because they contain ADP, or because they contain ATP/ATPγS but the side chain of the Arg^307^-finger residue, which is required for ATP hydrolysis, is disengaged.

|  | ClpX subunit |  |  |  |  |  |  |  |
| --- | --- | --- | --- | --- | --- | --- | --- | --- |
|  | A | B | C | D | E | F | substrate | ClpX |
| DHFR | ATP | ATP | ATP | ADP | ADP | ADP | DHFR-ssrA | $\Delta N$ |
| 8ET3 | ATPγS | ATPγS | ATPγS | ATPγS | ADP | ADP | SspB/GFP-ssrA | full-length |
| 6WRF | ATPγS | ATPγS | ATPγS | ATPγS | ATPγS | ADP | GFP-ssrA | $\Delta N$ |
| 6WSG | apo | ATPγS | ATPγS | ATPγS | ATPγS | ATPγS | GFP-ssrA | $\Delta N$ |
| 6VFS | ADP | ATP | ATP | ATP | ATP | ADP | GFP-ssrA | $\Delta N$ |
| 6VFX | ATP | ATP | ATP | ATP | ATP | ADP | GFP-ssrA | $\Delta N$ |
| 6PP5 | ATPγS | ATPγS | ATPγS | ATPγS | ATPgS | ADP | unknown | $\Delta N$ |
| 6PP6 | ATPγS | ATPγS | ATPγS | ATPγS | ATPγS | ADP | unknown | $\Delta N$ |
| 6PP7 | ADP | ATPγS | ATPγS | ATPγS | ATPγS | ATPγS | unknown | $\Delta N$ |
| 6PP8 | ATPγS | ATPγS | ATPγS | ATPγS | ATPγS | ADP | unknown | $\Delta N$ |
| 8E91 | ATPγS | ATPγS | ATPγS | ATPγS | ADP | ADP | none | full-length |
| 8E8Q | ATP | ATP | ATP | ATP | ADP | ADP | none | $\Delta N$ |
| 8E7V | ATP | ATP | ATP | ATP | ADP | ADP | none | $\Delta N$ |

**Figure S1.**
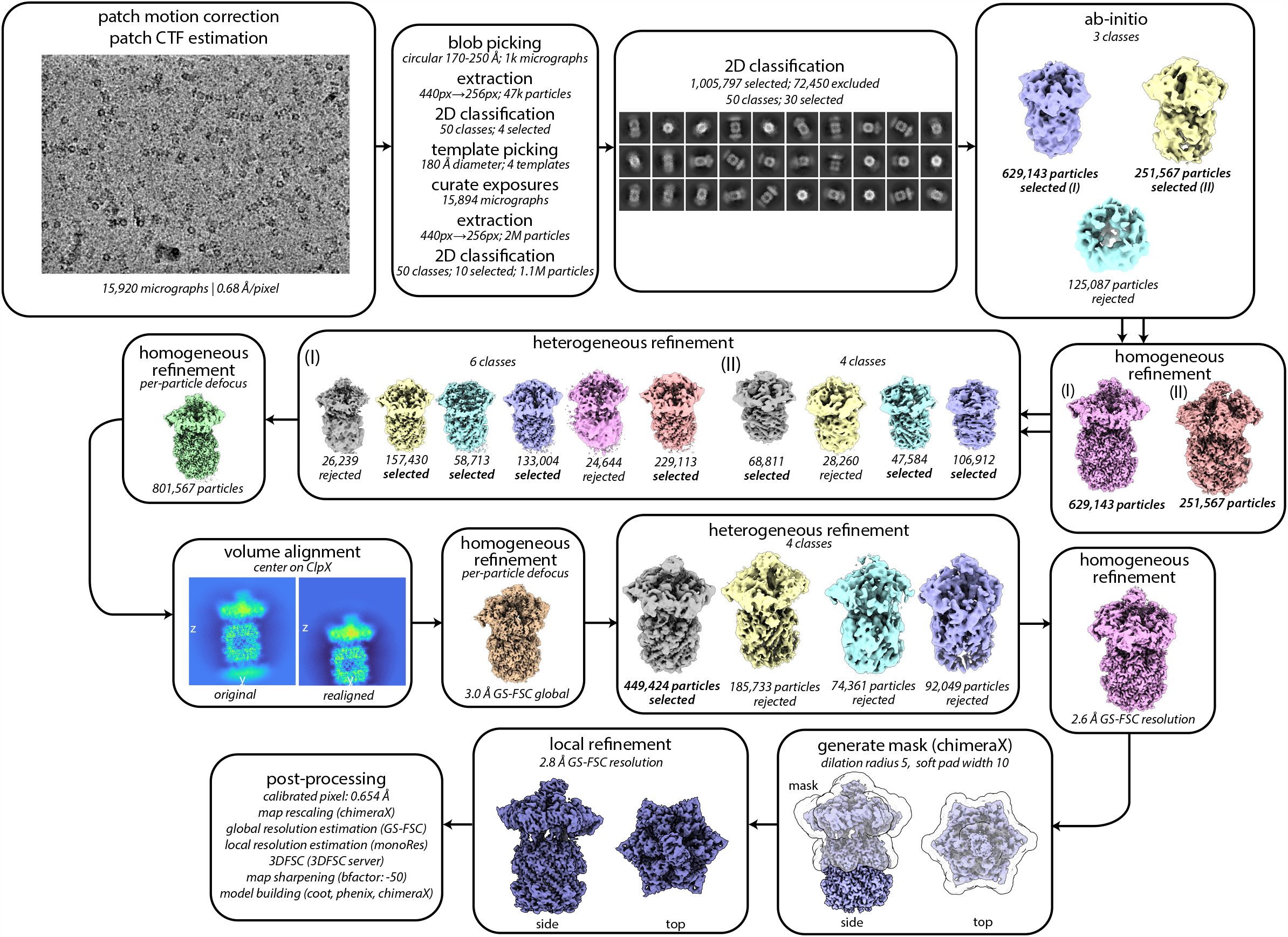
Image processing workflow. CryoSPARC processing workflow for single-chain ClpX^ΔN^/ClpP/DHFR•MTX particles. Job names, job details, and non-default parameters (italicized) are noted in each box.

**Figure S2.**
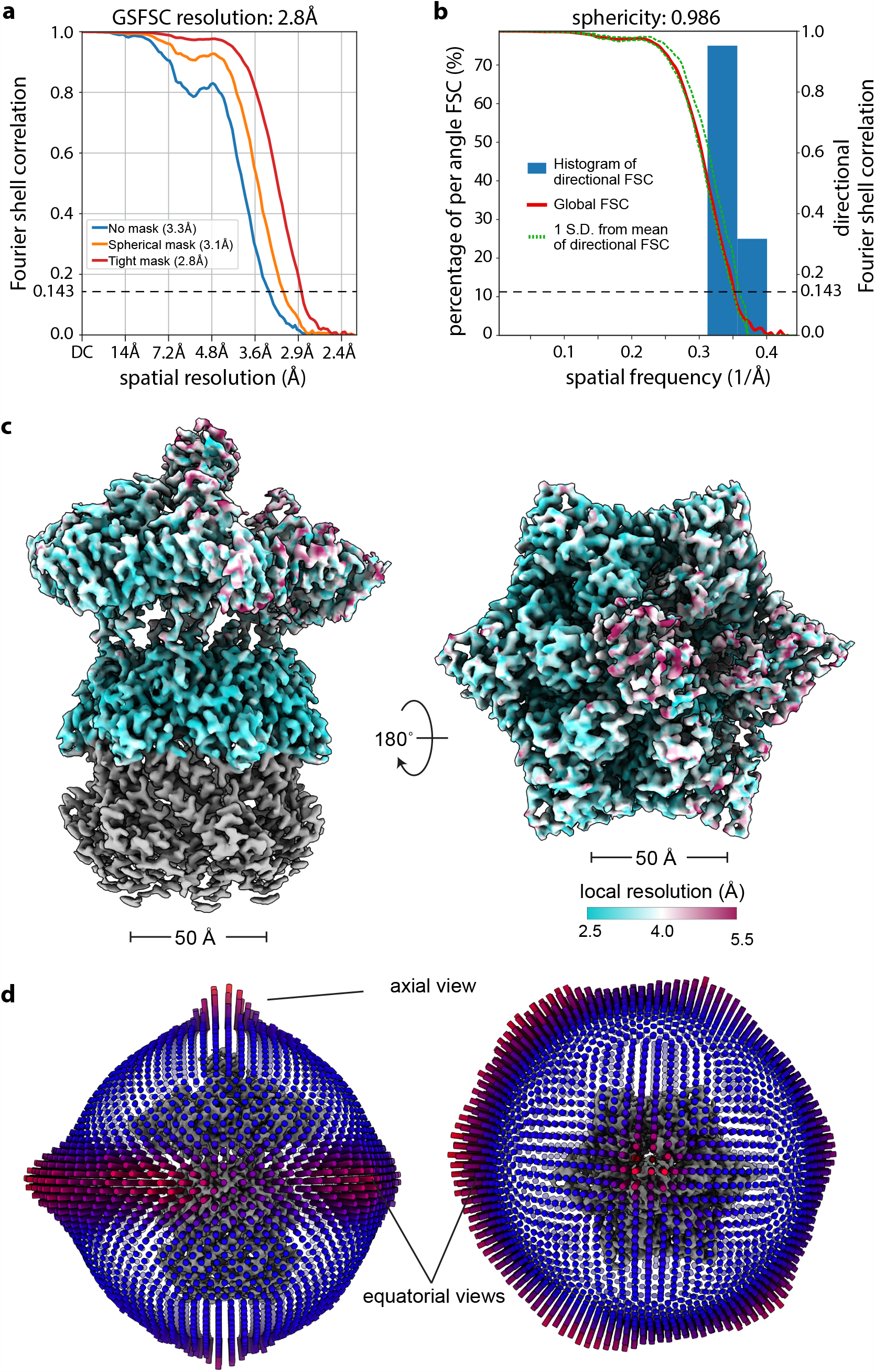
Estimates of GSFSC resolution. **(a)** Global resolution estimated by the gold-standard Fourier Shell Correlation method as implemented in CryoSPARC (Punjani, Rubinstein *et al*. 2017). **(b)** Directional FSC as estimated by the 3DFSC server (Tan, Baldwin *et al*. 2017). **(c)** Density map colored by local resolution as estimated by cryoSPARC’s implementation of monoRes (Vilas, Gomez-Blanco *et al*. 2018). Regions outside of the local refinement mask colored grey. **(d)** Projection-angle distribution.

**Figure S3.**
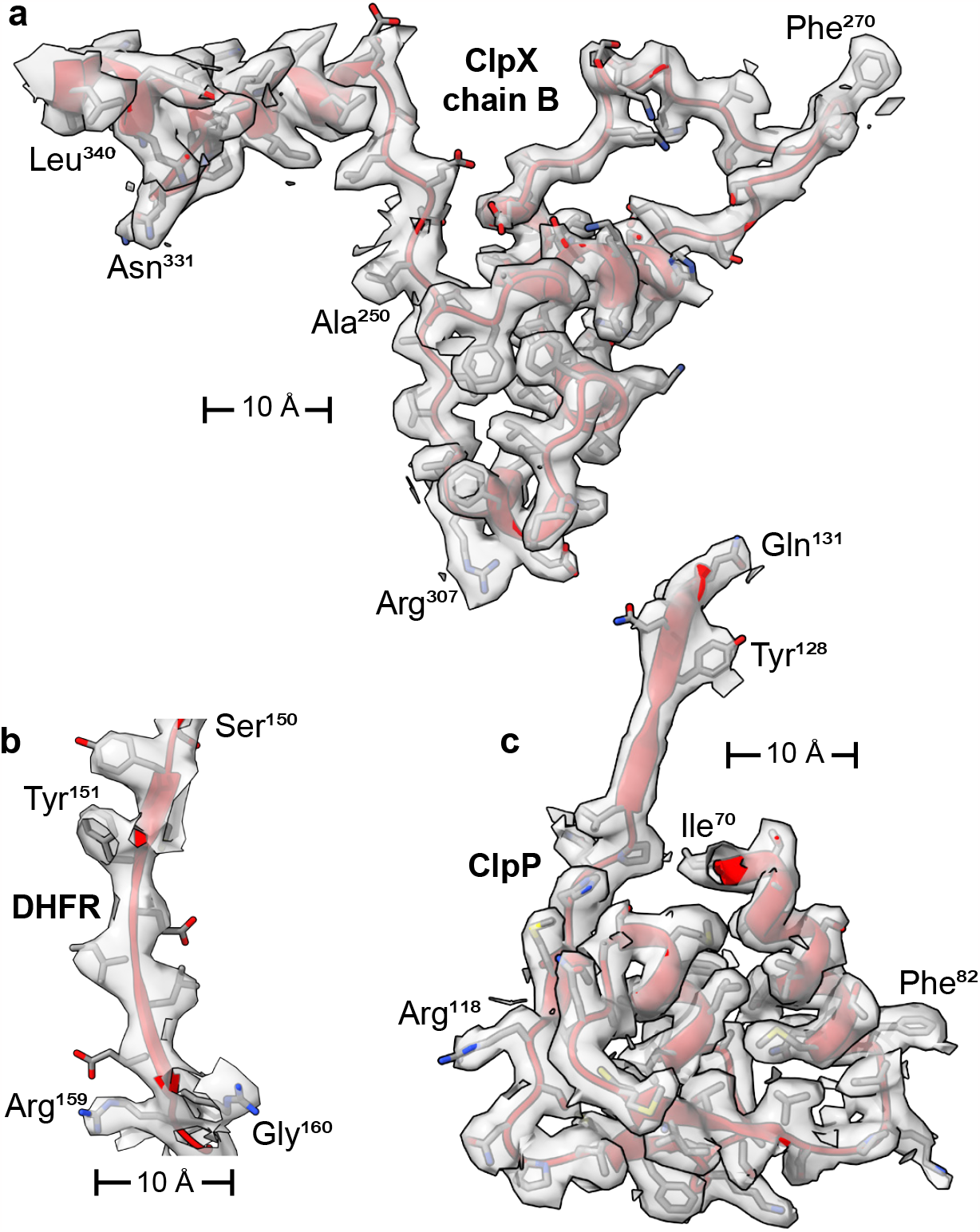
Cryo-EM density map and atomic model. Cryo-EM density map (grey semitransparent surface) overlayed on the fitted atomic models, with secondary structure elements colored red, and sidechains colored by atom type. **(a)** ClpX residues 270-340 of chain B. **(b)** DHFR residues 150-160. **(c)** ClpP residues 131-170 of chain i.

**Figure S4.**
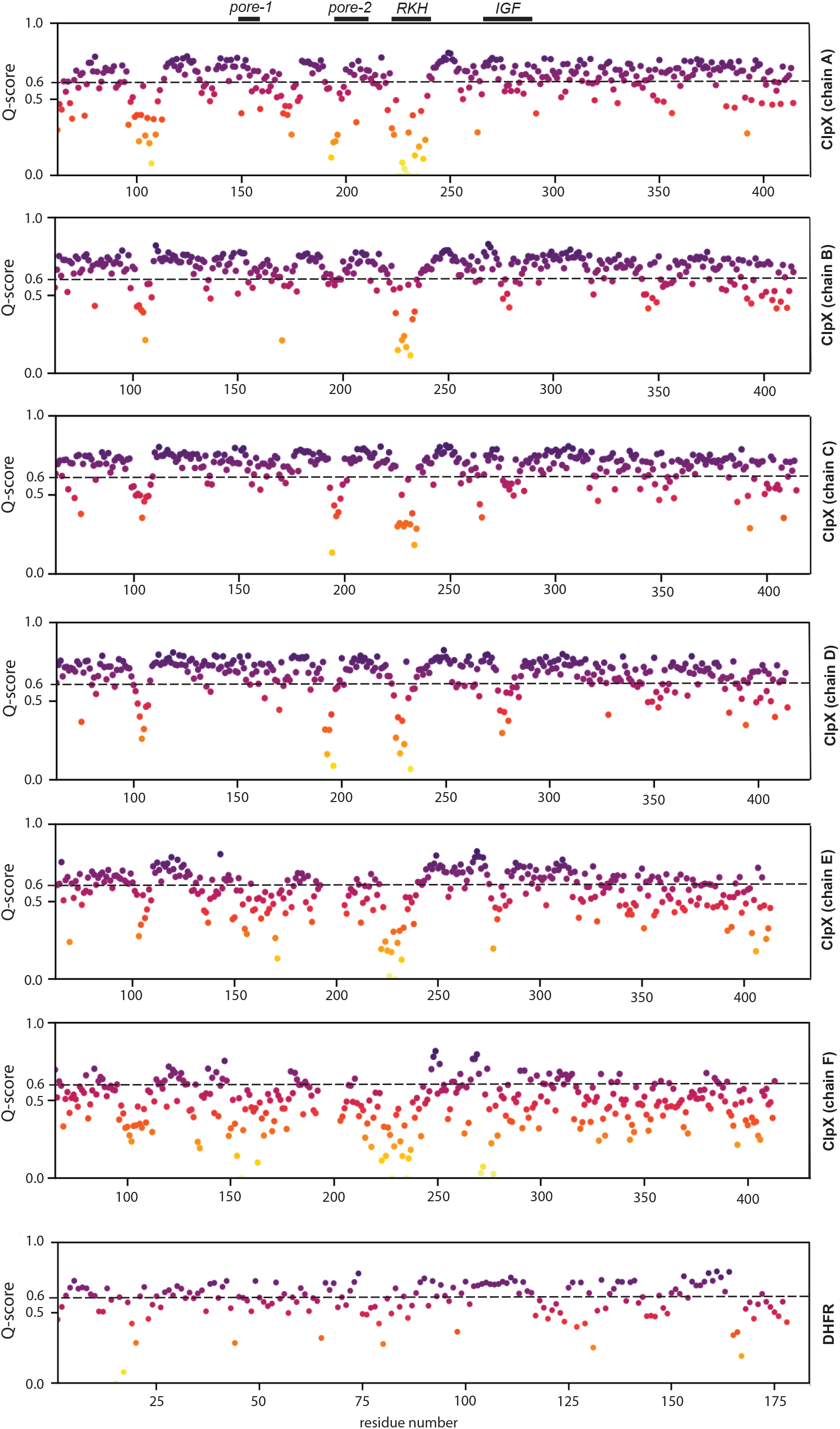
Map-model assessment. Calculated Q-scores (Pintilie, Zhang *et al*. 2020) for ClpX subunits and DHFR. Expected Q-score (0.6) given map resolution noted, and location of ClpX flexible loops annotated.

**Figure S5.**
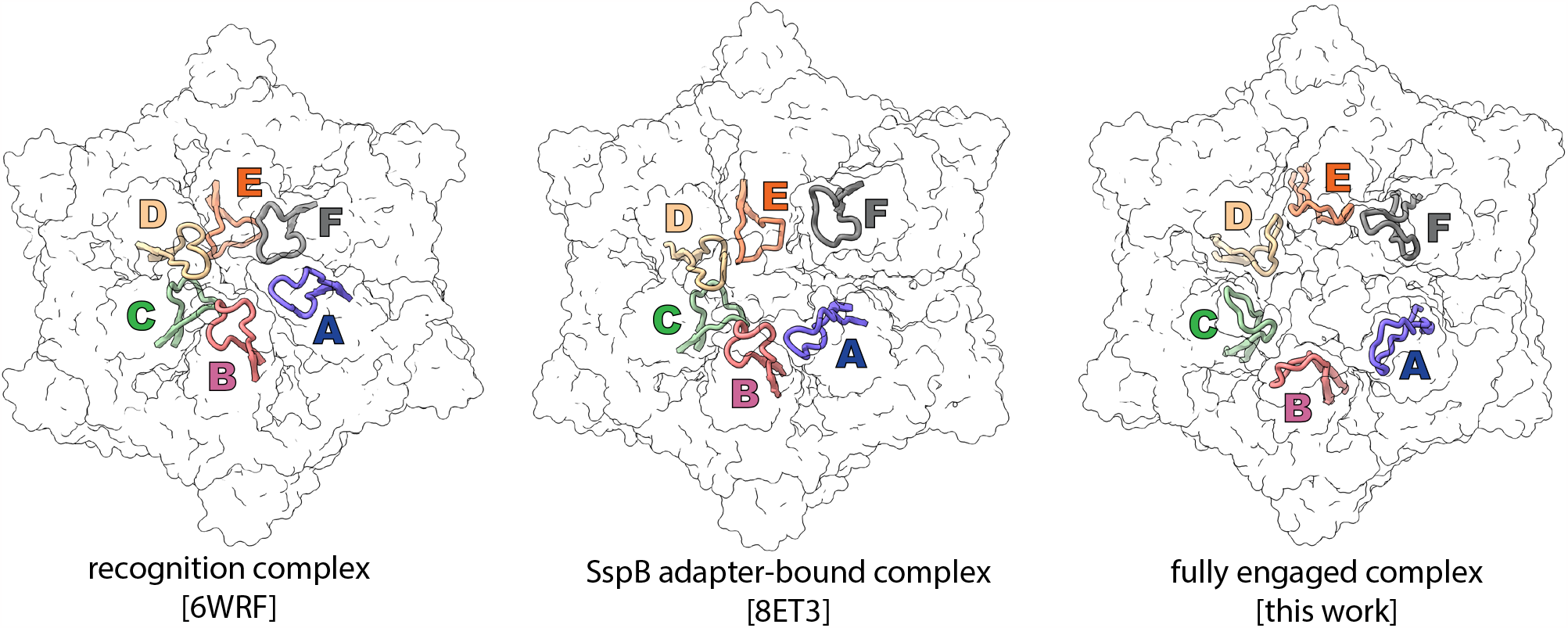
Conformational flexibility of the ClpX RKH loops. Diverse conformations of RKH loops (residues 218-240) from ClpXP structures 6WRF (left) (Fei, Bell *et al*. 2020), 8ET3 (center) (Ghanbarpour, Fei *et al*. 2023), and the DHFR-bound structure presented in this paper (right). Subunit colors: A (purple), B (salmon), C (green); D (wheat), E (orange), and F (gray).

**Figure S6.**
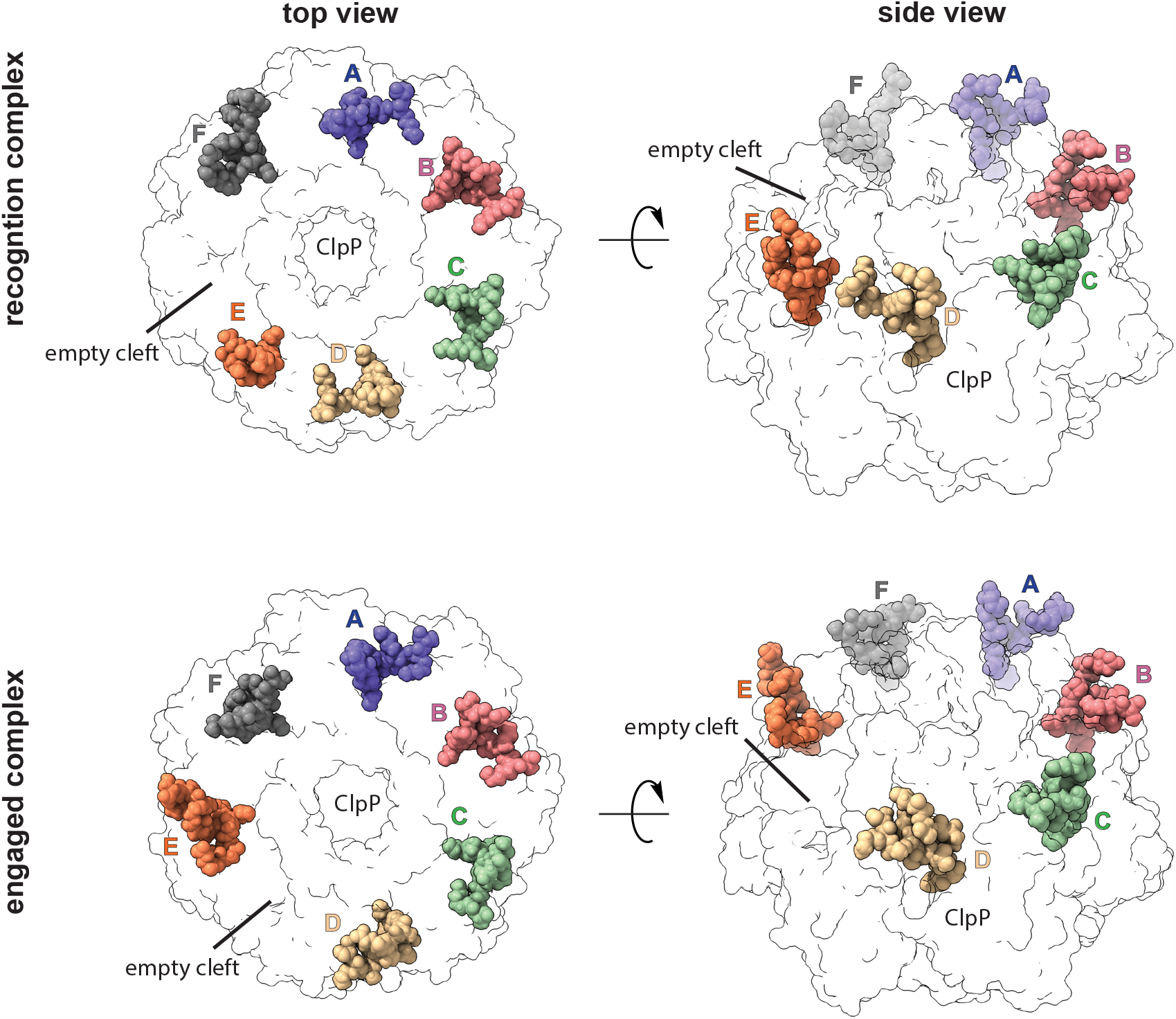
Rearrangement of ClpX-ClpP contacts. In a complex of ClpX bound to an ssrA degron (pdb 6WRF), the empty binding cleft on a ClpP heptamer is between the IGF loops of ClpX subunits E and F (top row). This arrangement is observed in most ClpXP structures (Fei, Bell *et al*. 2020, Fei, Bell *et al*. 2020, Ripstein, Vahidi *et al*. 2020, Ghanbarpour, Cohen *et al*. 2023). In the DHFR-engaged ClpXP structure (bottom row), the IGF loop of chain E moves into a binding cleft on the surface of the ClpP heptamer that is unoccupied in ClpXP structures except 8ET3 (Ghanbarpour, Fei *et al*. 2023). These loop docking interactions are depicted from top (left column), or side (right column) views.

**Figure S7.**
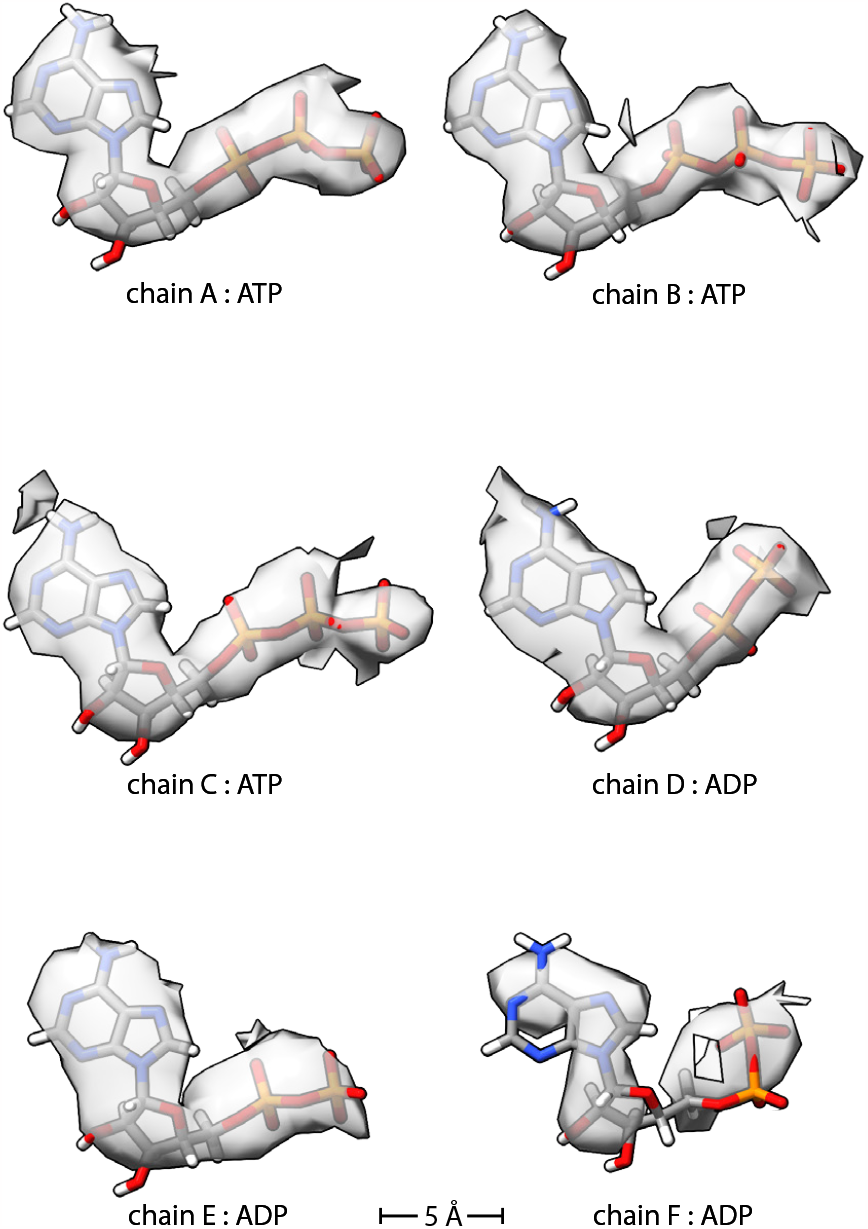
Density for ATP or ADP bound to different ClpX subunits in the DHFR-bound structure. Density map (grey semi-transparent surface) is overlayed on atomic models, which are colored by atom type.

**Figure S8.**
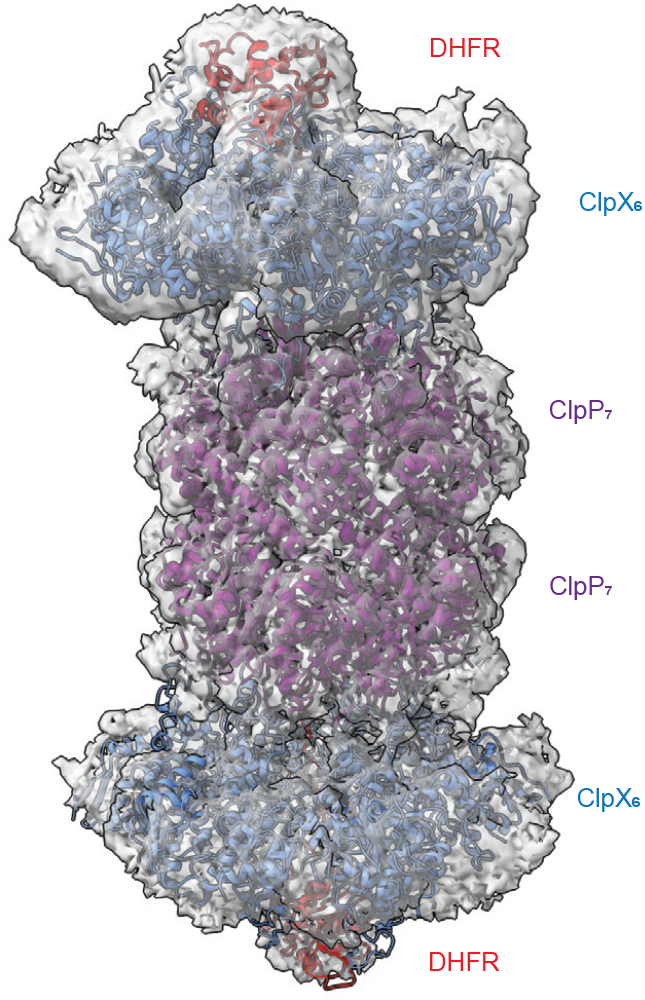
Low-resolution structure of a second ClpX•DHFR complex bound to the bottom heptameric ring of ClpP_14_. The distal ClpX•DHFR complex adopts multiple registers in relation to the top complex and has lower resolution as a consequence of conformational averaging.

## Notes

### Competing Interest Statement

The authors have declared no competing interest.

